# AI-Assisted Chemical Probe Discovery for the Understudied Calcium-Calmodulin Dependent Kinase, PNCK

**DOI:** 10.1101/2022.06.01.494277

**Authors:** Derek J. Essegian, Valery Chavez, Rabia Khurshid, Jaime R. Merchan, Stephan C. Schürer

## Abstract

PNCK, or CAMK1b, is an understudied kinase of the calcium-calmodulin dependent kinase family which recently has been identified as a marker of cancer progression and survival in several large-scale multi-omics studies. The biology of PNCK and its relation to oncogenesis has also begun to be elucidated, with data suggesting various roles in DNA damage response, cell cycle control, apoptosis and HIF-1-alpha related pathways. To further explore PNCK as a clinical target, potent small-molecule molecular probes must be developed. Currently, there are no targeted small molecule inhibitors in pre-clinical or clinical studies for the CAMK family. Additionally, there exists no experimentally derived crystal structure for PNCK. We herein report a three-pronged chemical probe discovery campaign which utilized homology modeling, machine learning, virtual screening and molecular dynamics to identify small molecules with low-micromolar potency against PNCK activity from commercially available compound libraries. We report the first described inhibitor hit series for PNCK that will serve as the starting point for future medicinal chemistry efforts for hit-to-lead optimization of potent chemical probes.

## Introduction

Understudied kinases, a designation assigned by the common-fund NIH project, Illuminating the Druggable Genome (IDG), comprise 24% of the entire human kinome (1-3). This number reflects our collective gap in knowledge and reveals biases in research towards certain classes or families of kinases. For instance, the CAMK family (Calcium Calmodulin Dependent Kinase) is understudied as a whole, with no targeted small molecule inhibitors to CAMKs in clinic or under clinical investigation despite an abundance of preclinical evidence suggesting pervasive roles of CAMKs in disease progression(4-11). Conversely, kinases of the TK (Tyrosine Kinase) family represent only 2% of the understudied kinases(1). Understudied kinases are denoted as such due to their lack of biological annotations, experimentally derived crystal structures, potent small molecule inhibitors and bibliographic references (3). Such dearth of information presents significant hurdles to the discovery of novel drug targets and subsequently novel small molecule therapeutics. However, analysis of large-scale multi-omics data has allowed for the identification and prioritization of understudied kinases whose expression and activity may correlate with disease activity or disease prognosis. Using data from The Cancer Genome Atlas (TCGA), we have identified several CAMKs, in particular, PNCK (CAMK1b), to be clinically relevant in several solid-tumor cancers(1, 12). Expression of PNCK was on average 6log2-fold greater in tumor samples compared to adjacent normal tissue in clear cell renal cell carcinoma (ccRCC) patients. Further, high expression correlated significantly with survival, tumor size and histological grade in all kidney cancer cohorts. To study the biological relevance of PNCK in vitro and validate this kinase as a novel target, an AI-assisted chemical probe discovery campaign was initiated.

Due to lack of chemical and biological data, machine learning models and homology models were generated to enrich libraries of over 7 million commercially available small-molecule compounds to test for inhibitory activity against PNCK.

## Materials and Methods

### Homology Modeling

Protein models and homology models were generated using Prime (Schrödinger Release 2019-4: Prime, Schrödinger, LLC, New York, NY, 2020). For protein models, crystal structures were imported into Prime by PDB ID (4FG7, 4FG8, 4FG9). Using Prime Protein Preparation Wizard, missing loops and side chains were filled in, missing hydrogens were added, and the protein underwent restrained energy minimization. Water molecules with less than 3 contacts with the protein were removed. For homology modeling of PNCK, the canonical amino acid sequence was obtained from UniProt (Q6P2M8). All homology models underwent loop refinement using serial loop sampling. For all models, extended molecular dynamic simulations (500ns-1us) were run to assess model validity and stability. All models contained co-crystallized ligands and as such, ligands were manually removed and then re-docked into the binding site to assess for model accuracy in predicting the experimentally derived binding pose and residue contacts and interactions. Homology models were assessed via Ramachandran plots to ascertain allowed and disallowed dihedral angles of residues in comparison to resolved parent crystal structures. Additionally, our generated models were compared to those predicted by Alpha Fold via structural alignment and RMSD (13).

### Docking Grid Generation

Receptor docking grids were generated for virtual screening with Glide Receptor Grid Generation (Schrödinger Release 2019-1: Glide, Schrödinger, LLC, New York, NY, 2019) and OpenEye’s Make_Receptor function (14) (OpenEye Scientific Software, Santa Fe, NM. http://www.eyesopen.com.) All proteins were initially prepared using Schrödinger Protein Prep Wizard prior to grid generation. Grids were then generated by using the co-crystallized ligand as the centroid for docking similarly sized ligands. Therefore, de novo grids in the active sites did not need to be created. Docking restrictions were made in the grid to ensure certain hydrogen-bond interactions were maintained during docking, particularly interactions with the hinge residues. Default settings were used in both OpenEye and Schrodinger grid generating modalities.

### Molecular Dynamics (MD) Analyses

All-atom explicit solvent Molecular Dynamic (MD) simulations were run using the Desmond GPU Accelerated suite of Schrodinger (**Schrödinger Release 2019-1** Desmond Molecular Dynamics System, D. E. Shaw Research, New York, NY, 2020. Maestro-Desmond Interoperability Tools, Schrödinger, New York, NY, 2020.) Simulation times varied between 300ns and 3μs, with a maximum of 3000 frames and a recording interval of 1/1000th of the overall simulation time. NPT (Isothermal-Isobaric) ensemble was used for the simulation, which most accurately represents laboratory conditions at ambient temperature and pressure (300K and 1.01325 bar). Each model system was relaxed before simulation for several nanoseconds. Systems were built also using Desmond with the SPC solvent model and OPLS3e forcefield. Simulations were computed across dedicated GPU cores.

### Machine Learning and Compound Enrichment

3 unique virtual screening campaigns were used to prioritize commercially available compounds to purchase for biochemical screening: A naïve Bayesian classifiers model, a neural net model, and a Tanimoto color and shape screen. Our group has previously developed ligand-based Laplacien-modified Naïve Bayesian classifiers to predict small molecule kinase activity trained on small molecule activity data across the entire human kinome(15). A list of homologous proteins was obtained via Clustal Omega multiple sequence alignment tool using the primary sequence of the PNCK kinase domain obtained from Uniprot(16, 17). The top 31 results had sequence identity homology which ranged from 70% to 30%, with annotated active compounds with IC50s <100nM (AURKA, AURKB, AURKC, BRSK1, CAMK2D, CAMK2G, CHEK2, DAPK3, FER, MAPKAPK2, MKNK2, MYLK, NUAK1, PDPK1, PLK4, PRKAA1, PRKACA, PRKACB, PRKACG, PRKD1, PRKD2, PRKD3, PRKG2, PRKX, RPS6KA1, RPS6KA2, RPS6KA3, RPS6KA5, SGK1, STK17A, TSSK1B). Amongst selected kinases, there were 9,693 active small molecule compounds in the training set. The model was then tested against over **7 million** commercially available compounds from the eMolecules library. Compound predictions from eMolecules were then scored based on their probability of binding to said target kinase-using a score known as EstPGood. A probability cut-off of 0.1 was used to enrich the virtual screening library. Additionally, compounds with multiple target predictions were prioritized while those with only one target prediction were filtered out. Compounds with a molecular weight <250 g/mol or >700 g/mol were also filtered. In total, 49,512 unique compounds were prioritized for the first virtual screen.

A second machine learning effort, a neural net-based approach, was also used which compiled data from the KKB (Kinase Knowledge Base)(18) and the eMolecules library of commercially available compounds. For the neural net machine learning predictions, 314 kinases with 331,985 known active compounds were used for building the model. Compounds were featurized with Deepchem and ECFP4 fingerprints with a size of 1024 bits. The dataset was split randomly into 10% test, 10% verification and 80% training. There were 4 layers in a multiclass tensorflow network with the first layer being the 1024-bit input layer, the second layer having 2000 nodes, the third layer having 500 and the final layer being the output layer of 314 kinases. Prediction results came from the eMolecules library of compounds that were then predicted for their activity against this network and stored for later retrieval. The same homologous kinases used in the Naïve Bayesian model were used to retrieve predictions from the neural net. 289,008 unique compounds were identified as predictions to be used in the virtual screen. Compounds were filtered with a probability cutoff of >0.1, compounds active against >1 kinases, and with molecular weights ranging from 250-700g/mol.

A third approach took advantage of the binding poses of ATP in the active site of PNCK homology models. A Tanimoto shape and color screen was employed using OpeneEye’s fastROCS GPU implementation. Compounds were screened in Enamine, MolportDB and eMolecules libraries. Libraries were prepared in OpenEye first using QUACPAC which involves tautomer and protomer enumeration at physiological pH ranges. OMEGA was then used to generate up to 150 conformers for each ligand. The eMolecules dataset was filtered before the shape screen using the OpenEye “ Lead-Like” filter, generating 1,169,867 unique compounds. For Enamine, the advanced HTS screening library was used which contained 97,410 unique compounds while MolportDB Screening Compound Library had 287,773 compounds. A Tanimoto combination score cutoff of 0.5 (which is the sum of a color score and shape score) was used to prioritize compounds for docking using HYBRID.

### Ligand Library Preparation

Compounds selected for virtual screen were populated as SMILES in a comma-separated file along with a unique identifier or catalog number obtained from the vendor. Using Pipeline Pilot (BIOVIA **Pipeline Pilot**, Release 2018, San Diego: **Dassault Systèmes)**, SMILES were canonicalized and the file was transformed to a .SDF structure file to be used as input for LigPrep (**Schrödinger Release 2019-3**: LigPrep, Schrödinger, LLC, New York, NY, 2019) or for OpenEye QUACPAC and OMEGA (OpenEye Scientific Software, Santa Fe, NM. http://www.eyesopen.com). For large-scale virtual screens with over 100,000 compounds, OpenEye Hybrid docking was utilized. Following enrichment by Hybrid Docking Score, smaller subsets of ligands were then used to dock using Glide (Schrodinger) for further binding pose visualization and MD simulations. Ligand libraries were obtained from Enamine’s Advanced HTS library, MolportDB and eMolecules. The Enamine library was curated to have compounds with molecular weights between 300 and 500g/mol. The MolPortDB had no molecular weight filters, with compounds ranging from 150 to 750 g/mol. The eMolecule library was filtered into screening subsets using both lead-like and drug-like filters provided by OpenEye because the library was too large to screen as a whole. For ligand preparation, default settings were used in both programs. Compounds to be used in OpenEye were prepared first using QUACPAC which involves tautomer and protomer enumeration. OMEGA was then used to generate up to 150 conformers for each ligand. For GLIDE docking, ligands were prepared using LigPrep (**Schrödinger Release 2021-1**: LigPrep, Schrödinger, LLC, New York, NY, 2021) which generates all conformers, tautomers and ionization states at physiologic pH ranges.

### Ligand Shape Screen

Shape screens were conducted using fastROCS GPU (OpenEye Scientific Software, Santa Fe, NM. http://www.eyesopen.com), which can screen millions of compounds in seconds. Structure files of ATP exported from docking models of PNCK were used as input for the screens. A combined Tanimoto shape and color score was used to filter the most similar compounds in eMolecules, MolportDB and Enamine prepared compound libraries.

### Ligand Docking

For large scale docking studies utilizing over 100,000 structures, OpenEye’s HYBRID Dock was used. HYBRID docking was done in parallel on all PNCK grids in one job thus aggregation of scores was not necessary as HYBRID only reports the highest scoring pose per compound per grid. The top 25,000 docking poses were exported for final analysis. For smaller scale docking jobs or confirmatory docking studies prior to MD simulation, Glide Docking was used (**Schrödinger Release 2019-1**: Glide, Schrödinger, LLC, New York, NY, 2019). All default settings were used. Glide standard precision docking (SP) was performed using 3D ligands generated from LigPrep. Up to 25 poses were generated per ligand with the top 5 poses being selected for output. Glide scores were exported for aggregation across various grids using Pipeline Pilot.

### Compound Storage and Cataloging

All compounds were purchased as powders in quantities less than 5mg. Compounds were dissolved in various organic solvents (Methanol, Dichloromethane, ethyl acetate) and were aliquoted out in separate vials. Solvents were evaporated using a rotovap and high vacuum and aliquoted powders were stored at −20C. Compounds were dissolved in DMSO to a concentration of 10mM prior to use in assays. Compounds were stored in DMSO at −80C no longer than 1 week. All purchased compounds were catalogued virtually using the Collaborative Drug Discovery (CDD) Vault (https://www.collaborativedrug.com/).

### Biochemical PNCK Inhibitor Screen

ADP-Glo Kinase Assay (**V6930, Promega)** was used to screen the first round of compounds against PNCK. GST-tagged PNCK recombinant protein was purchased from Novus (NBP1-99886). Autocamtide-II (GenScript RP10271) was used as the peptide substrate. This peptide was dissolved in water and prepared immediately prior to the assay. Calcium/Calmodulin 10X (Abcam ab189137) solution was used to create a custom kinase buffer: 40mM Tris,pH 7.5; 20mM MgCl2; 0.1mg/ml BSA; 50μM DTT and Ca2+/Calmodulin solution (0.03μg/μl, Calmodulin, 1mM Tris,pH 7.3, 0.5mM CaCl2). The assay was run according to protocol using 25uM of ultra-pure ATP. A standard curve was generated per run which displayed the linear range of luminescence at various ratios of ATP to ADP. A standard curve was then done to determine the amount of protein needed to generate a signal in the linear range of the ATP curve. 100ng of PNCK and 2ng of substrate was used per well in the 96-well plate format of this assay. Compounds were screened at 10uM and incubated at RT for 1-2h. Plates were read using the Clariostar Plus. Data was normalized to a DMSO and staurosporine controls.

### Cell-based PNCK inhibitor Screen

The NanoBRET Target Engagement Intracellular Kinase Assay (Promega **N2520)** is a cell-based binding screen which reads loss of BRET signal as a result of target binding. A PNCK Nano-luciferase fusion vector plasmid was generated custom by Promega. The K-10 tracer was used for the assay, as determined by Promega. The plasmid was maxiprepped by Genewiz for use in all future assays. HEK293 cells were seeded in a T175 flask in DMEM with 10% FBS. The following day, cells were counted to 10,000, resuspended in assay media (Optimem with no phenol red, 1% FBS). Plasmid and transfection reagent (Fugene HD, Promega **E2311**) were added to the mixture and cells were seeded in a white 96-well plate. After 20-24 hours, media was changed, and the assay proceeded according to protocol. K10 tracer was added at the recommended concentration of 1uM while compounds were initially screened at both 10uM and 1uM. Cells incubated with compound at 37 degrees Celsius for a maximum of 2 hours before the plate was read using Clariostar Plus. Data was normalized to Staurosporine and DMSO controls.

### Data Analysis

Data analysis of ADP Kinase Glo results were carried out per the manufacturer’s instructions. Results were represented as a percent decrease in activity. Data was normalized to staurosporine (positive control) as 100% decrease and DMSO (negative control) as a 0% decrease. Data for NanoBRET was analyzed by dividing the acceptor by the donor emission and dividing by 1000 to obtain raw millibret units (mBU). Background was subtracted using control emission averages. Data was normalized to positive and negative controls of staurosporine and DMSO. Concentration-response curves were plotted in GraphPad Prism 8 and IC50s were extrapolated using non-linear fit with concentrations transformed to logarithmic scale. Baseline was held constant and represented the average mBU of staurosporine controls while the top was constrained to represent the average mBU of DMSO controls. Error bars on dose-response curves represent standard deviations.

## Results

### Building and Assessing PNCK Homology Models

No experimentally derived crystal structure exists for PNCK. However, several related structures in the CAMK family have been resolved which could be used as a structural template to generate a homology model. To determine the CAMK member with greatest homology to PNCK, the amino acid sequence of PNCK (isoform 1) was used in pBLAST (NCBI). PNCK canonical amino acid sequence was obtained from Uniprot(16) (ID: Q6P2M8) and the kinase domain is predicted to be from residues 15-270. CAMK1a was identified as the most homologous structure in the PDB with sequence identity of >70% in the kinase domain(19). Three individual iterations of CAMK1a structures were selected as templates. All structures contained an N-terminal lobe of 5 anti-parallel beta-sheets and the canonical alpha-helix with a C-terminal lobe composed of primarily alpha helices. All native CAMKs contain a C-terminal region with auto-inhibitory and CaM binding sequences, assuming a helix-loop-helix structure(6, 19-22). CAMKIa (1-320) (PDB: 4FG9, 2.40 Å) represents a full-length version of the calmodulin kinase with the regulatory auto-inhibitory domain intact. Therefore, this kinase was experimentally shown to be inactive despite being co-crystallized with ATP in the active site. PDB structures 4FG8 (1-305, 2.20 Å) and 4FG7 (1-293, 2.70 Å) are also co-crystallized with ATP but represent two distinct forms of the kinase. 4FG8 is a truncated structure with no calmodulin binding site, thus it is in an inactive state. 4FG7 has the entire regulatory domain removed and thus represents a constitutively active form. It has been determined that the regulatory domain of CAMK1a inhibits kinase activity by interacting with the N-terminal lobe and masks the Thr177 phosphorylation site, restraining the kinase in an inactive conformation(19). The regulatory domain, however, does not occlude nucleotide binding site, thus 4FG9 (1-320) is able to bind ATP. The activation loop still adopts an inactive conformation, with a “ DFG-up” like state whereby the phenylalanine is not completely rotated out of the binding pocket, representative of a distinct inactive conformation(19, 23). 4FG8 (1-315) similarly adopts an inactive conformation, with no magnesium detected in the active site. 4FG7 (1-293) displays the structural elements of a kinase in active conformation, particularly the alpha-helix displays interactions with its conserved glutamate, forming salt bridges with ATP. This structure has a true, active, “ DFG-in” conformation. In all three ATP-bound structures, the activation loop is largely disordered; contrasted with the Apo-structure, whereby the activation segment adopts a helical formation. The use of three unique templates of CAMK1a allowed for a broad sampling of conformational states in the virtual screen.

Each homology model was created using PRIME and included the entire protein with ATP in the binding site (**See Methods**) (**Figure 1A-C)**. Loops were extensively refined, and the structure went through multiple rounds of energy minimization to produce the most stable model. To determine if the model is valid, the co-crystallized ATP was removed from the binding site and was docked back in using Glide. A model was deemed valid if it accurately predicted the pose and binding mode which was present in the experimentally derived structure within 1-2 Angstroms. The PDB structures all have a conserved interaction with the adenine amino group and a glutamate or valine in the hinge. Additionally, there are hydrogen bonds between the catalytic lysine (Lys44) to the gamma phosphate group and hydroxyl groups on the sugar. For PNCK, the predicted hinge residues included Val93 and Glu91 with the gatekeeper residue being Met90 and the catalytic lysine at Lys44. Models were finally evaluated for stability through extended all-atom explicit water molecular dynamic (MD) simulations **(Supplemental Figure 1)**. MD simulations were run using Desmond (GPU implementation) for 500ns (**see Methods**). An analysis of the RMSD of the C-alpha backbone as a function of time was used to assess overall structure integrity; as the change of RMSD approaches zero, the structure “ converges” to a local stable conformation. Each PNCK model converges at about 150-300ns to an average RMSD of about 3-5 angstroms (**Supplemental Figure 1**). Models were additionally assessed using SwissModel Structure Assessment with mol prob scores >2 and Ramachandran allowed residue percentages greater than 90% **(Supplemental Figure 2, Table 1)**. Finally, structures were compared to those predicted by Alpha Fold via structural alignment and RMSD (**Table 2**). To further increase the conformational space sampled during docking, the trajectories of the 500ns molecular dynamic simulations were clustered and 5 structurally diverse poses for each of the three homology models were extracted (**Figure 1D**). Thus, in total, 15 unique representations of PNCK were used in parallel for the virtual screening docking campaign.

**Figure 1:**
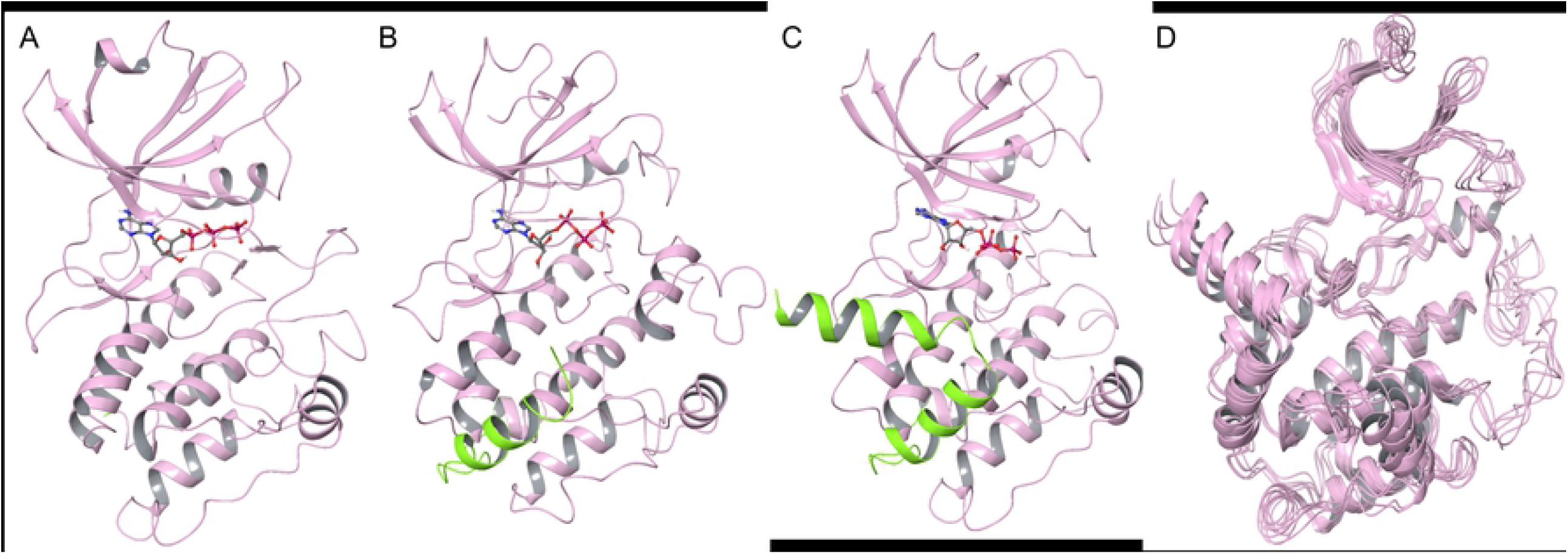
PNCK Homology Models with ATP Docked. A) 4FG7 Homology model with no CAM-binding site or auto-inhibitory domain. B) 4FG8 Homology Model with CAM-binding site (Green helix) and no autoinhibitory domain. C) 4FG9 Homology model with full length PNCK including autoinhibitory domain and CAM-binding site (Green helix). D) Overlay of 5 representative poses extracted from the 500ns molecular dynamics (MD) simulation trajectory of the 4FG9 homology model.

**Table 1:** SwissModel Structure Assessment.

| Model | MolProbity Score | Ramachandran Favored % | Ramachandran Outliers % |
| --- | --- | --- | --- |
| **4FG7-Homology** | 1.39 | 93.31% | 0.00% |
| **4FG7** | 2.07 | 95.87% | 0.00% |
| **4FG8-Homology** | 1.35 | 92.44% | 0.69% |
| **4FG8** | 2.31 | 95.38% | 0.00% |
| **4FG9-Homology** | 1.66 | 93.38% | 0.66% |
| **4FG9** | 2.05 | 96.88% | 0.00% |

**Table 2:**
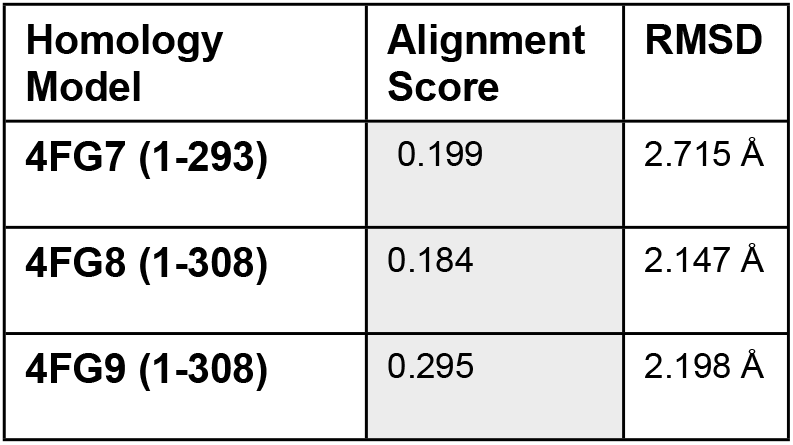
Structural Alignment of Homology Models using AlphaFold.

| Homology Model | Alignment Score | RMSD |
| --- | --- | --- |
| **4FG7 (1-293)** | 0.199 | 2.715 Å |
| **4FG8 (1-308)** | 0.184 | 2.147 Å |
| **4FG9 (1-308)** | 0.295 | 2.198 Å |

### Curating Compound Libraries for Scalable Ligand-based Virtual screening

In total, three unique virtual screening campaigns were used to prioritize commercially available compounds for *in vitro* screening. Each combined a scalable ligand-based approach with docking and molecular dynamics. Specifically, we used Naïve Bayesian classification models, a multi-task deep neural network model, and 3D shape-based screening (**see Methods**).

Our group has previously developed ligand-based Laplacien-modified Naïve Bayesian classifiers to predict small molecule kinase activity. These models have also been used to predict novel small molecule inhibitors of EGFR and BRD4(15, 24). The predictors were trained on small molecule activity data across the entire human kinome. Although there are no published active molecules against PNCK, similarity amongst kinase binding sites and cross-kinase activity of many kinase inhibitors facilitate the enrichment of PNCK inhibitors using predictive models of the most similar kinases. A list of homologous kinase domains was obtained by multiple sequence alignment with Clustal Omega using the primary sequence of PNCK kinase domain obtained from Uniprot(16, 17) (**See Methods)**. Most of the homologous kinases with sufficient small molecule data for high-quality predictors belonged to the CAMK family, including DAPK3 and BRSK1; other kinases included PRKACA, AURKA and AURKB. The model was then tested against over 7 million commercially available compounds from the eMolecules library. In total, 44,736 unique compounds were prioritized for the first virtual screen.

More recently, we developed a kinome-wide multi-task deep neural network classifier trained on aggregated curated data from ChEMBL(25) and the KKB (Kinase Knowledge Base)(18). Overall, 331,985 known active compounds across 314 kinases were used for building this model. Predictions were made for the entire eMolecules library. Similar to the previous approach, compounds predicted active for homologous kinases were prioritized (**See Methods**). Aggregating the predictions, a total of 289,008 unique compounds were prioritized. PRKACA was the kinase that was most represented, with 13,780 compounds predicted active.

A third approach took advantage of the three-dimensional predicted binding poses of ATP in each of the active sites of the PNCK homology models. A Tanimoto shape and color screen was employed using OpeneEye’s fastROCS (GPU implementation). Compounds from the Enamine, MolDB and eMolecules libraries were screened after generating 3D conformers. ROCS (Rapid Overlay of Chemical Structures) is a powerful method for shape similarity and pharmacophore screening which uses Gaussian overlay functions to measure both shape and “ color” similarity between compounds [180, 181]. ROCS utilizes both Tanimoto and Tvsersky functions to score overlap similarities. Volume overlap generates a shape score while color Tanimoto scores are derived from scoring similarities between electrostatic forces for functional atoms or chemical groups. The eMolecules dataset was filtered before the shape screen using the OpenEye “ Lead-Like” filter, generating 1,169,867 unique compounds. For Enamine, the advanced HTS screening library was used which contained 97,410 unique compounds while MolportDB had 281,987 compounds. A Tanimoto combination score cutoff of 0.5 (which is the sum of a color score and shape score), was used to prioritize compounds for docking using HYBRID. 163,493 compounds with Tanimoto scores above 0.5 were docked. The top 25,000 scoring compounds from each dataset were then evaluated further.

### Virtual Screening Campaign 1: Naïve Bayesian Classifiers and Docking

Compounds prioritized using the Laplacien modified Naïve Bayes models were prepared for docking using Schrodinger LigPrep. Compounds were then docked using Glide(26) into each of the 15 docking grids generated from the 3 homology models. Docking scores were then aggregated across all 15 receptors (**See Methods**). The top Glide Scores and mean Glide scores for each compound were used to rank and prioritize. As a reference, ATP received a glide average glide score of −8.59 kcal/mol. Docking scores for the machine-learning predicted compounds had an average of −5.52 kcal/mol and a standard deviation of −0.83. The top 1% of docking scores scored below (better than) −8.26 kcal/mol (**Figure 2)**. Compounds were first clustered using extended connectivity fingerprint methods with a radius of 6 (ECFP_6). Clusters were then ranked using the average glide score and top clusters were evaluated individually based on synthetic tractability of the scaffold, likely binding pose, and ability for derivatization. One to two compounds from each cluster were then selected for purchase to test in in *in vitro* screening. In total, 25 compounds were purchased from the eMolecules catalogue **(Supplemental Table 1)**.

**Figure 2:**
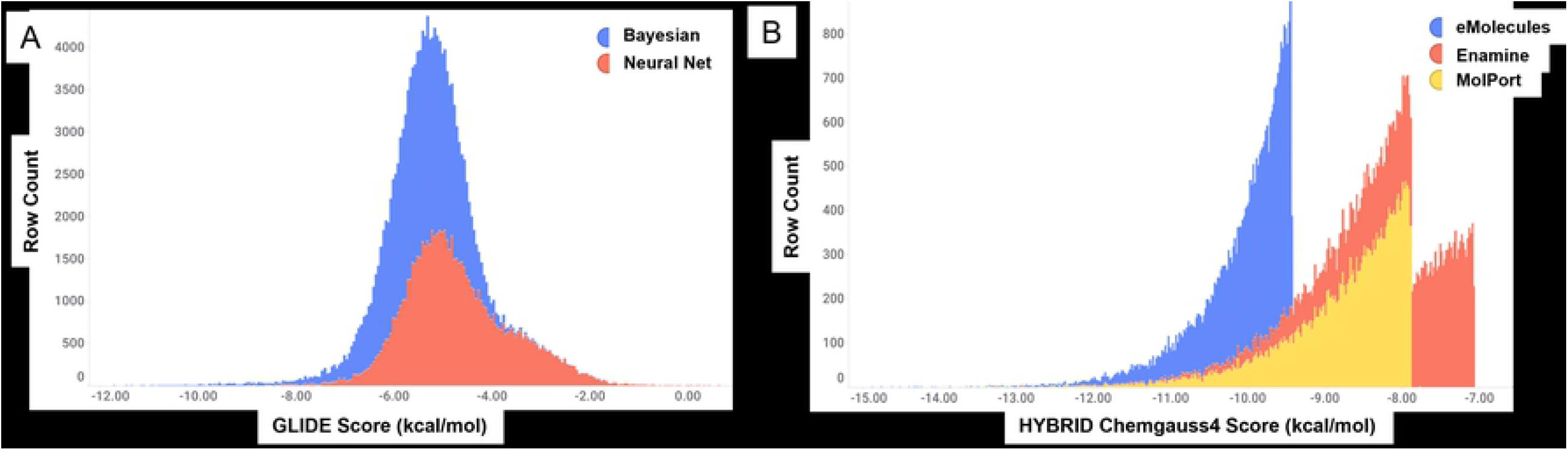
Distribution of Docking Scores in Different Machine Learning Models. A) The naïve bayesian model and neural net model were both used to screen for predictions against kinases homologous to PNCK in the eMolecules library. The distribution of docking scores was very similar suggesting the neural net did not offer many new predictions. This was confirmed by manual inspection of clustered scaffolds B) vROCs shape screening and HYBRID docking using OpenEYE was performed using three small molecule libraries. The composition of each library offered much more diversity and subsequently was reflected in the distribution of docking scores, with eMolecules compounds scoring the best.

Among compounds selected for biochemical screening, the docking scores ranged from −7.37 kcal/mol (UM_027) to −10.99 kcal/mol (UM_039) and molecular weights ranged from 315 (UM_019) to 410 (UM_027) g/mol. All compounds passed The REOS filter, (rapid elimination of swill) and PAIN filter screening (pan-assay interference) except for UM_036, as the presence of a 1,3-diazinane-2,4,6-trione may react with reagents used in such screens. Compounds UM_027, UM_029, UM_031, UM_034, and UM_039, all had at least one “ Rule of Three” violation, while none of the compounds were predicted to be exceptionally cell-permeable (Exceptional Cell permeability has a QPPCaCo score >500) (27). In fact, 4 compounds are predicted to be considered poorly cell permeable (QPPCaCo <25) (UM_021, UM_031, UM_034 and UM_039) from QikProp analysis (**Supplemental Table 2)**. Thus, compounds from this cohort were more “ drug like” (MW <500, cLogP <5, Molar Refractivity between 40-130) than “ lead like”.

### Virtual Screen Campaign 2: MTDNN (Multi-task Deep Neural Net) and Docking

As described above, MTDNN predictions were obtained using the same homologous kinases as campaign and 289,008 unique compounds were identified as predictions to be used in the virtual screen. Compounds were filtered with a probability cutoff of >0.5, activity to >1 kinase and with molecular weights ranging from 250-700g/mol leading to a final docking library of 39,787 compounds. Although PRKACA was the most overrepresented kinase, distributions of docking scores amongst the various kinase predictions did not vary significantly. The overall distribution of docking scores was slightly different compared to the naïve Bayesian classifier model, with the average docking scores shifting to −4.70 kcal/mol, with the best docking score being −9.69 kcal/mol **(Figure 2)**. In fact, many of the same scaffolds as in campaign 1 were similarly identified in this method. For example, the cyclopropyl-pyrazole-carbonyl moiety in UM_019 and UM_034 was present in many of the top clusters of the MTDNN campaign. Additionally, the thieno[2,3-d]pyrimidine scaffold found in UM_027, UM_028, UM_033 and UM_035 was also highly represented in these predictions campaign. Therefore, this method did not greatly increase the diversity of scaffolds, likely due to the fact that compounds were screened again the same eMolecules library and the two machine learning methods did not identity significantly different compounds. Therefore, no compounds identified from this method were purchased for further analysis in biochemical or cell-based screens.

### Virtual Screen Campaign 3: Tanimoto ROCS Shape Screen and Docking

By far, the greatest and most diverse docking results came from a third approach which simply consisted of finding molecules that are of similar shape, size, and electronic configuration as our computational model of ATP docked in the PNCK active site. Using fastROCS GPU of OpenEye, millions of compounds from eMolecules, Enamine and MolDB were queried against the 15 poses of ATP obtained from the molecular dynamic trajectories of the 3 PNCK homology models (**See Methods**). All conformers and ionization states generated from the compounds with Tanimoto combo scores >0.5 were then docked across all 15 receptors using OpenEye Hybrid and the top 25,000 scoring representations across all docking models were selected for further evaluation. Compounds were clustered using ECFP6 fingerprints as described before and top scoring clusters were evaluated for synthetic tractability and binding mode. As with the Naïve-Bayesian campaign, representative structures with unique scaffolds from the top scoring clusters were purchased for analysis in in vitro screens. Significant differences were noted in distribution of Tanimoto and docking scores amongst the three databases. Compounds from the eMolecules database had an average HYBRID docking score of −9.9 kcal/mol, followed by MolPortDB with an average of −8.56 kcal/mol and Enamine with an average of −7.92 kcal/mol (**Figure 2**).

When comparing the three homology models for PNCK, docking results from the Tanimoto screen using HYBRID show a strong preference for the 4FG9 grid (the inactive form of PNCK), with the best docking scores from these compounds coming from poses of the auto-inhibited state. Conversely, 42% of compounds selected from the Naïve Bayesian cohort (UM_019-UM_039) scored best with GLIDE in the 4FG7, constitutively active PNCK model.

## Target Engagement Assays

The 25 Compounds prioritized by Naive Bayesian classifiers and docking (Campaign 1) were initially tested in a cell-free biochemical kinase activity screen, ADP Kinase Glo (**See Methods**). Six compounds were identified as hits (>50% inhibition of kinase activity at 10uM): UM_026, UM_029, UM_032, UM_035, UM_038 and UM_039 (**Supplemental Figure 3)**. However, it was determined that PNCK is not an active kinase and has sub-optimal activity without phosphorylation at Thr177. Thus, significant amounts of PNCK (>100ng per reaction) were needed to obtain signals in the linear range of the standard curve to extrapolate kinase inhibitory activity. Therefore, NanoBRET target engagement assay was used for future screens, including re-testing all compounds tested in the ADP-Glo Kinase Screen. NanoBRET is more sensitive than the ADP-Glo and additionally selects for compounds that are cell permeable (**see Methods**).

From the shape screen and docking campaign, compounds UM_194 to UM_244 were prioritized for testing in the NanoBRET target engagement assay. All compounds chosen for binding assays were analyzed for their putative binding mode by GLIDE docking and molecular dynamics (MD) simulations using the homology model for which it scored the highest. 55 compounds were initially tested in the NanoBRET assay, including compounds UM_019-UM_039 from the first campaign. Compounds UM_192, UM_193 UM_242, UM_243 and UM_244 were not cell permeable while UM_234 and UM_245 had poor aqueous solubility, which was properly indicated by QikProp predictions. Additionally, UM_234 and UM_243 were flagged by the PAINS filter due to the presence of a thiophene structure and a catechol structure, respectively. As such, these compounds were not subject to further analyses. Compounds from each campaign were similar in physiochemical properties but compounds from the Tanimoto shape screen were significantly more cell-permeable (**Table 3)**. The NanoBRET screen was performed using two concentrations for compounds: 10µM and 1µM. Data was normalized to DMSO and staurosporine controls (**Figure 3**). Hits were defined as having greater than 50% inhibitory activity at 10µM. This assay confirmed actives and inactives that were identified via ADP kinase GLO with the exception of UM_037. UM_037 was only shown to inhibit ∼31% of PNCK activity at 10uM in the ADP GLO assay. However, in the NanoBRET screen, UM_037 was shown to be a potent binder at 10uM with 56% inhibition. Conversely, top compounds in the ADP GLO assay failed in NanoBRET, likely due to poor cell-permeability. In particular, UM_039 had 91% inhibition of kinase activity in the biochemical screen with negligible activity in the cell-based screen. QPPCaco predictors from QikProp (Schrodinger) score UM_039 at 8.374, which is predicts poor cell permeability.

**Table 3:**
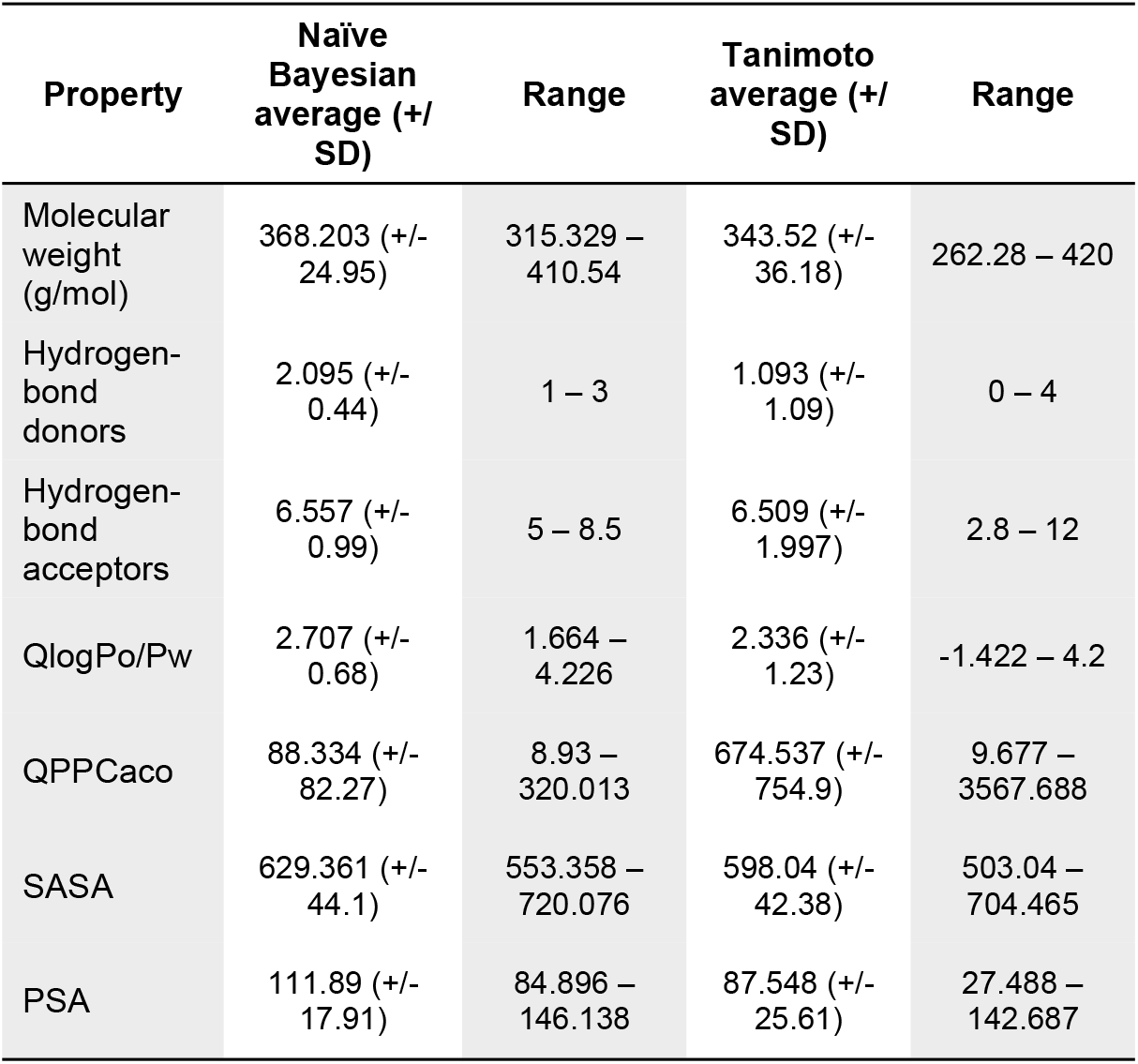
Physiochemical Properties of Compounds From Each Virtual Campaign.

**Figure 3:**
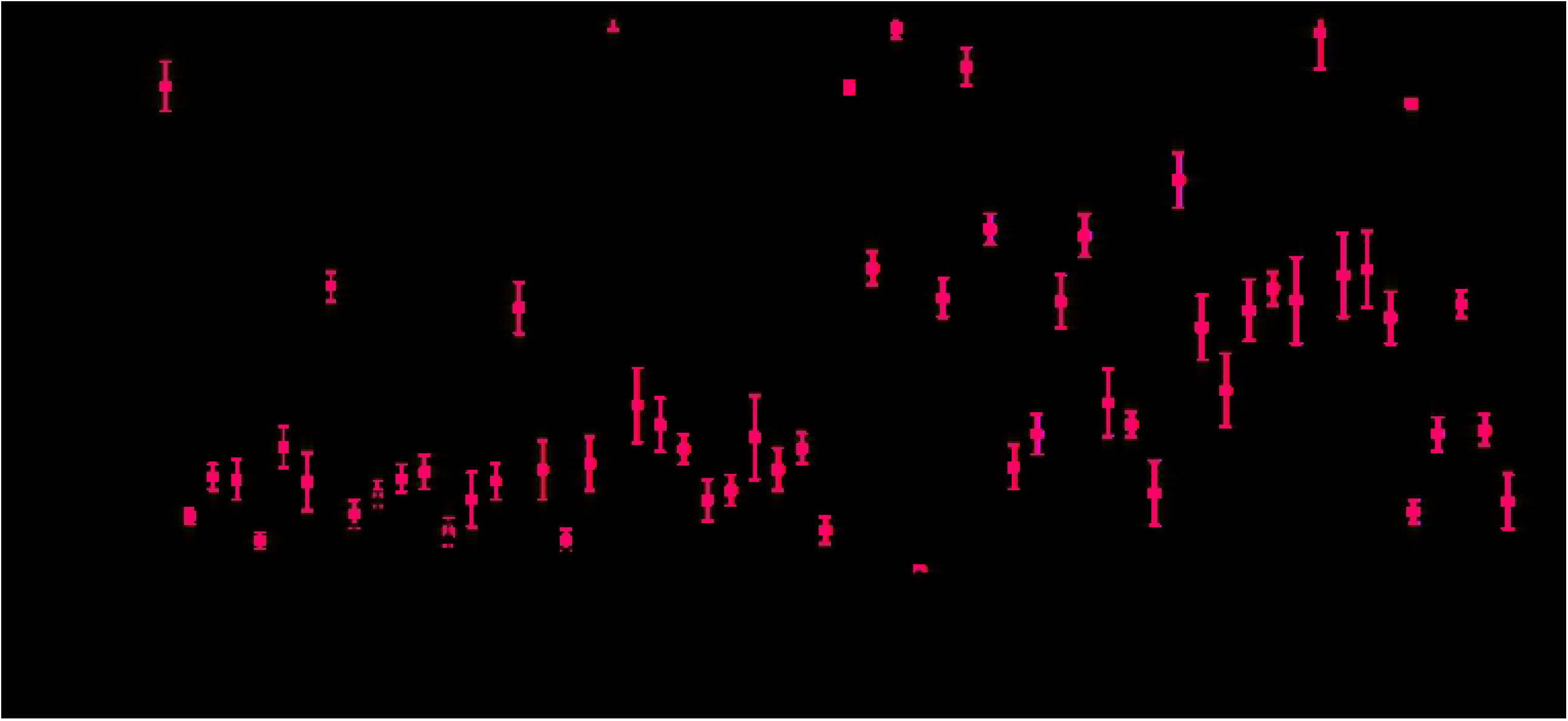
Cell-Based Screening of Prioritized Compounds. All compounds from the naïve Bayesian model in addition to compounds identified in the vROCs shape screen were tested in the cell-based target engagement assay, NanoBRET. Compounds were tested at concentrations of 10µM and 1µM. Activity was reported as normalized percent inhibition using DMSO and staurosporine as controls. Pan-CAMK inhibitor KN-62 was also used as a reference. Hits were identified as compounds with at least 50% inhibition at 10uM.

The top compounds chosen for follow-up validation in concentration-response curves were selected based on normalized percent inhibition at 10µM and 1µM, synthetic tractability and kinase inhibitor scaffold diversity or novelty (**UM_037, UM_195, UM_210, UM_213 & UM_228**). Concentration-response curves were generated for each compound using the NanoBRET system at 8 concentrations ranging from 24µM to 0.175µM. Nonlinear curve fitting after log-transforming of the concentrations allowed for calculation of IC50s (**Figure 4, Table 4)**.

**Figure 4:**
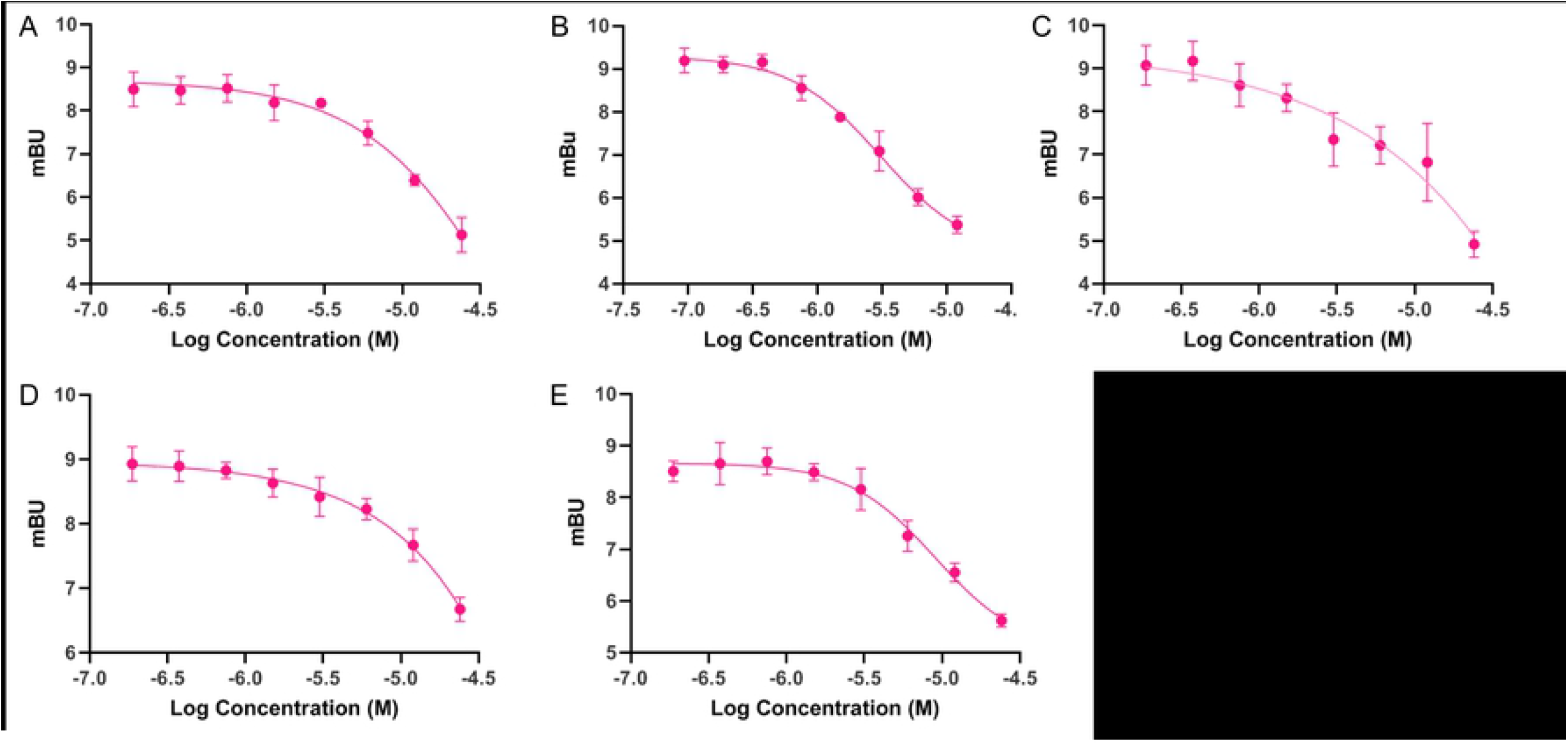
Dose-Response Curves for Top 5 Hit Compounds. A) UM_37, B) UM_195, C) UM_210, D) UM_213, E) UM_228

**Table 4:**
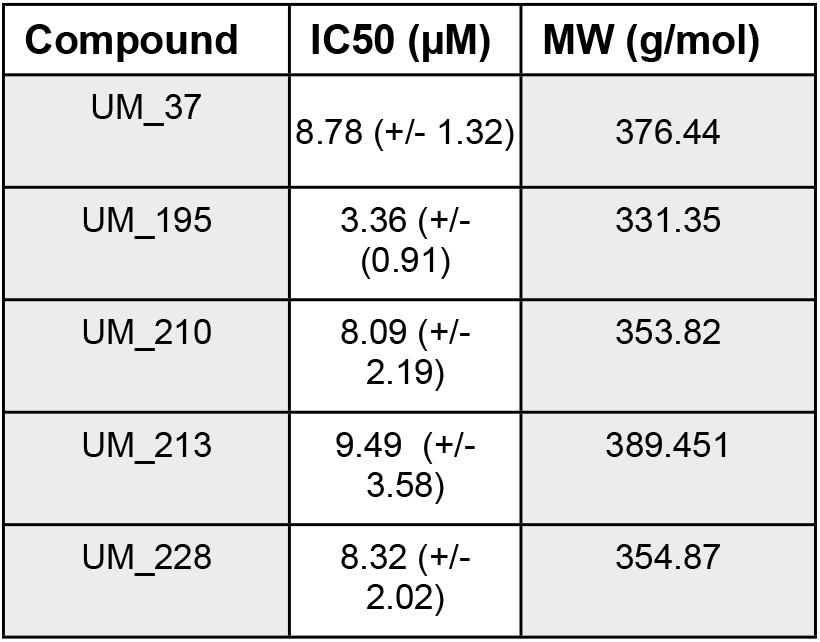
IC50s of Top 5 Hit Compounds.

| Compound | IC50 (μM) | MW (g/mol) |
| --- | --- | --- |
| UM_37 | 8.78 (+/- 1.32) | 376.44 |
| UM_195 | 3.36 (+/- (0.91)) | 331.35 |
| UM_210 | 8.09 (+/- 2.19) | 353.82 |
| UM_213 | 9.49 (+/- 3.58) | 389.451 |
| UM_228 | 8.32 (+/- 2.02) | 354.87 |

In conclusion, one top scaffold was identified from Naïve-Bayesian Classifier / docking campaign while the other 4 were identified through the Tanimoto nucleotide ligand shape and color screen & docking.

### Structure Activity Relationship (SAR) – Analogue by Catalogue

Each hit compound was studied again extensively to develop the most accurate binding hypothesis based on classical binding poses kinase inhibitors. Using the most likely predictive pose from initial docking studies **(Figure 5)** and MD simulations (**Supplemental Figures 4-23**), analogs were identified to potentially maximize hydrogen bonding interactions in the binding pocket, particularly at the hinge residues and to gain evidence for its true binding mode. Parent hit compounds were queried in ChemSPACE to find analogs among billions of chemicals in “ REAL space” that had variation at one or more R-group sites (**Supplemental Figures 4-23)** (28). Analogs were then docked in the PNCK models to analyze any improvement in docking scores based off additional hydrogen bond and hydrophobic interactions at the hinge, gatekeeper, or catalytic lysine residues. Analog compounds with improvement in docking scores or with modifications congruent with a structure-activity relationship hypothesis were ordered for synthesis on demand. Several custom analogs were ordered for each initial hit (limited by commercial availability) and were tested again in the NanoBRET screen at 1µM and 10µM (**Figure 6)**. Compounds with at least 50% activity at 10µM were analyzed further with concentration-response curves to generate IC50 values (**Figure 7**). These data then allowed for preliminary structure-activity relationships for each of the scaffolds.

**Figure 5:**
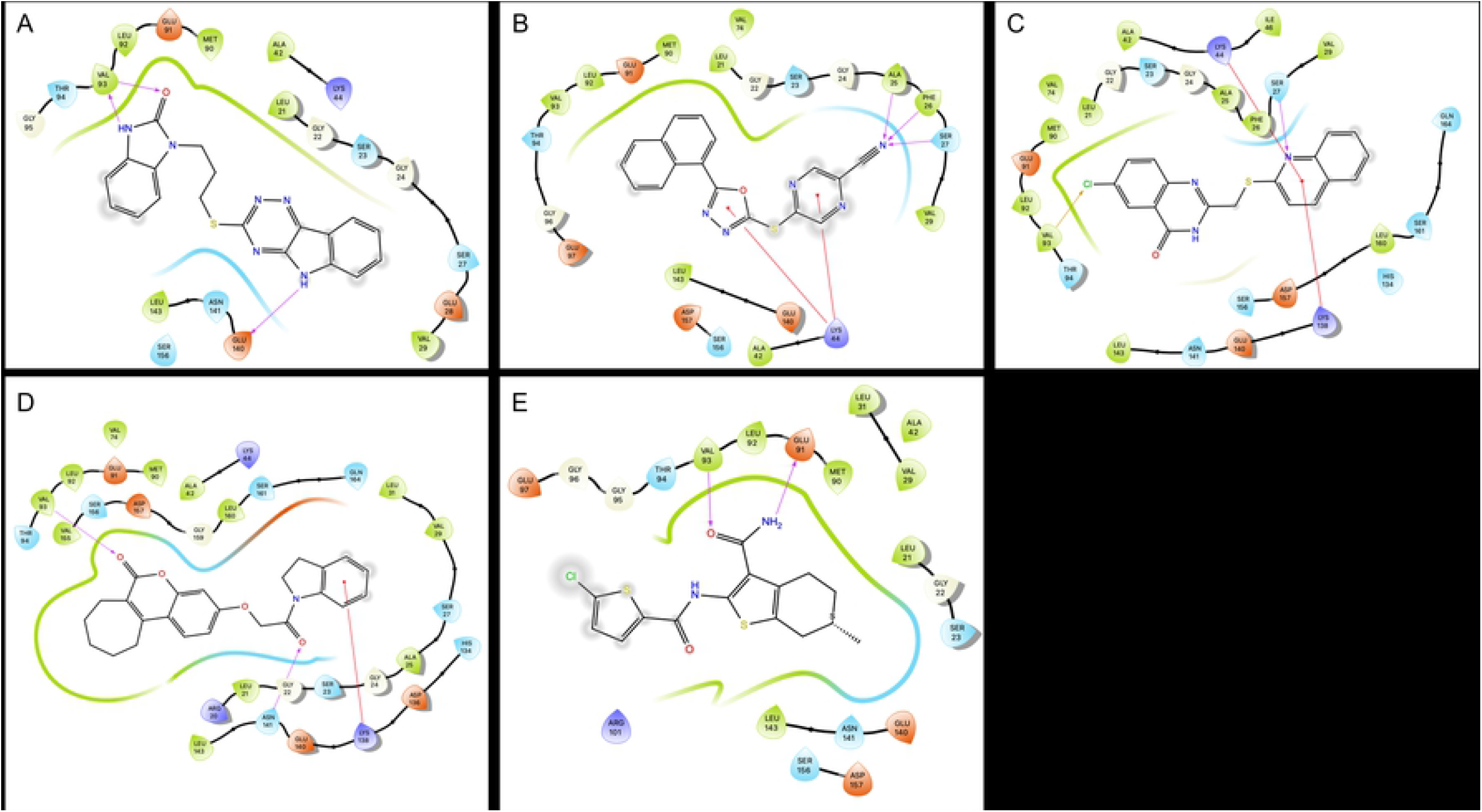
2D Representation of Predicted Binding Poses of Hit Compounds. A) UM_37, B) UM_195, C) UM_210, D) UM_213, E) UM_228

**Figure 6:**
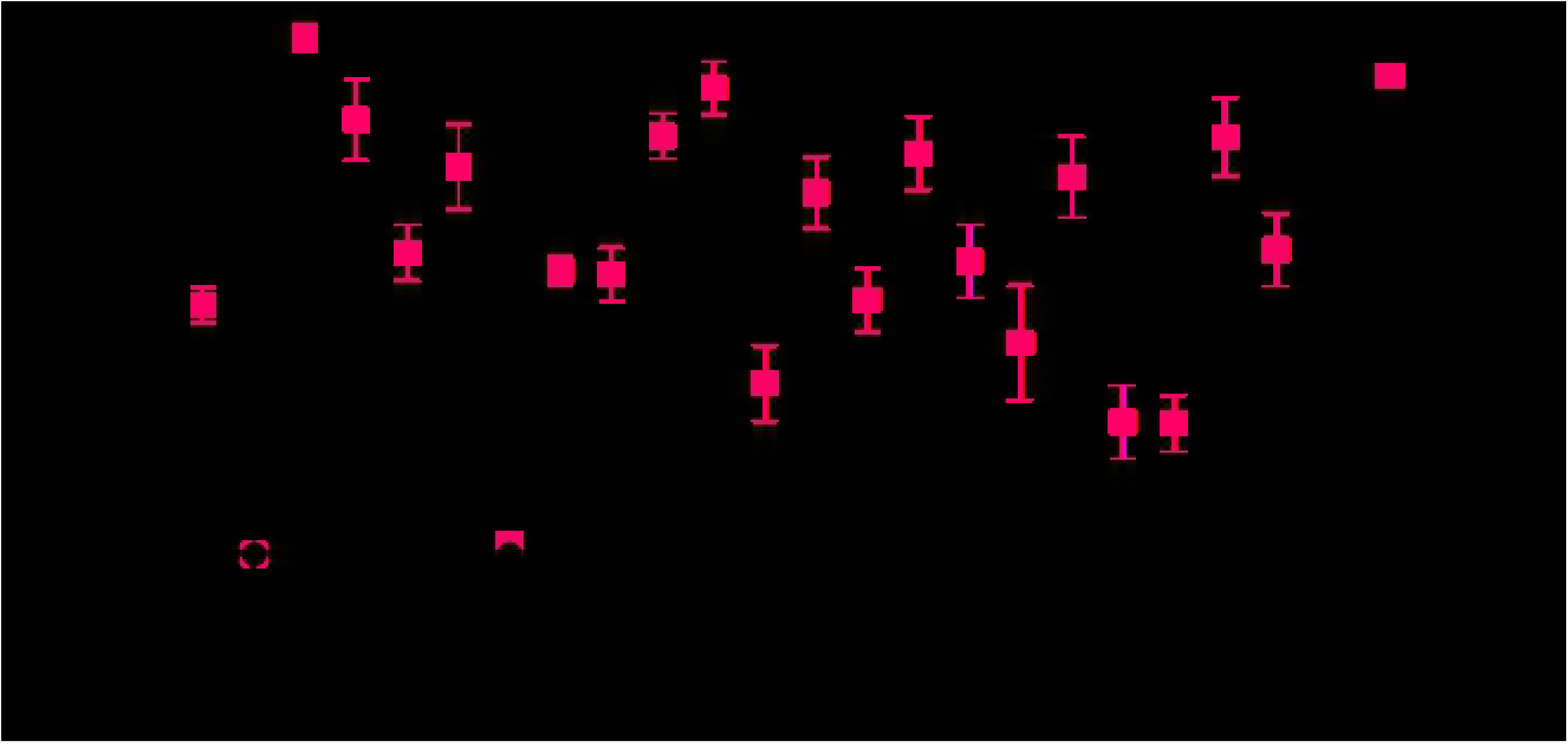
Cell-Based Screening of Analog Compounds: Analogs of the 5 hit compounds were identified from ChemSpace and prioritized via docking and SAR hypothesis. Compounds were tested using the NanoBRET assay at concentrations of 10 µM and 1 µM.

**Figure 7:**
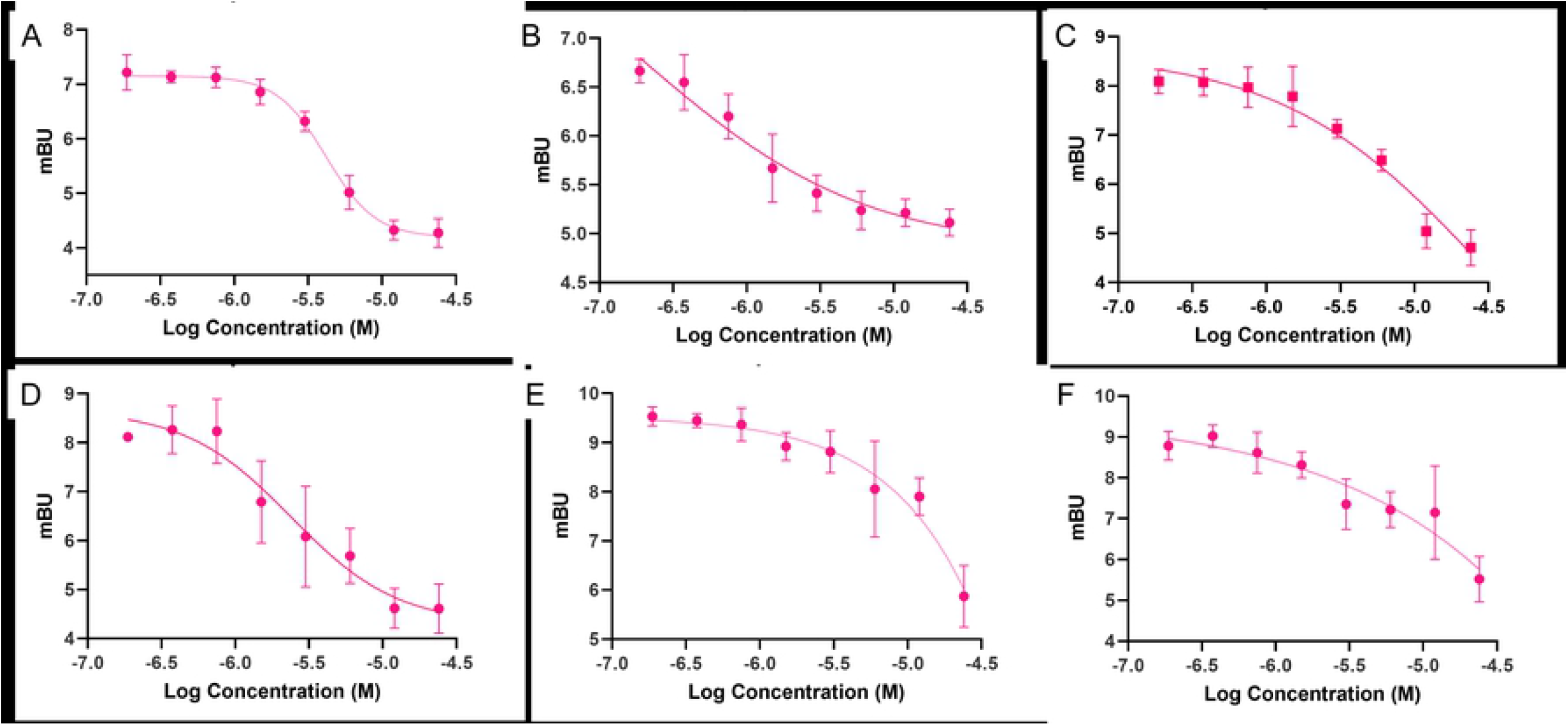
Dose Response Curves of Top Analog Compounds. A) UM_251, B) UM_252, C) UM_254, D) UM_259, E) UM_277, F) UM_280

**UM_228**,

(6R)-2-(5-chlorothiophene-2-amido)-6-methyl-4,5,6,7-tetrahydro-1-benzothiophene-3-carboxamide: Molecular modeling of UM_228 led to several binding hypotheses. As we prioritized poses with hydrogen bonds at the hinge, the predicted pose with the chlorine atom solvent exposed, the thiophene group deep in the binding pocket and the amide group positioned for hydrogen bonding with hinge residues Glu91 and Val93 was used for selecting analogs. Additional poses reversed the orientation by 180 degrees with the methyl group being solvent exposed and the chlorine participating in bonding interactions at the catalytic lysine. In selecting analogs, various parts of the molecule were considered to determine which groups of the scaffold were important for binding, determined by analyzing docking studies and MD simulations (**Supplemental Figures 4-6**). The methyl group was removed in one analog or replaced with bulkier groups to maximize hydrophobic interactions in the hinge. Chlorine was removed or was replaced with various hydrogen bond donors or acceptors such as a nitro group or various heterocycles. Docking studies of the analogs supported the hypothesis that the chlorine-substituted thiophene was likely situated towards the hinge, as replacing this with a purine group as in UM_251 greatly improved docking scores with sustained interactions with Val93 (**Supplemental Figure 7)**. Molecular dynamics analyses of UM_251 in the 4FG9 model demonstrates various hydrogen bond interactions along with significant mobility of the auto-inhibitory domain to create a new binding pocket, with new interactions gained at Serine156 (**Supplemental Figure 8**). In UM_252, thiophene in UM_228 is replaced with a fluorophenyl-pyrazole. Binding hypotheses situate the fluorophenyl group in the hinge with sustained hydrophobic interactions during molecular dynamic simulations while the pyrazole and amide groups participated in multiple hydrogen bonds and water bridges to Val93 and Ser156; more than the parent compound 228 (**Supplemental Figure 9)**. This was supported via NanoBRET assays with analogs UM_251 and UM_252 improving from the IC50 of UM_228 by one order of magnitude (**Figure 6**).

**UM_213**,3-[2-(2,3-dihydro-1H-indol-1-yl)-2-oxoethoxy]-6H,7H,8H,9H,10H,11H-cyclohepta [c]chromen-6-one: All 3 docking models oriented UM_213 into the binding pocket of PNCK with the pyrone carbonyl hydrogen bonding with Val93 and the cycloheptane ring pointing towards either the solvent or towards to the gatekeeper, Met90. Additionally, there were hydrogen bonds between Lys44 and Ser156 with the amide nitrogen and oxygen. Molecular dynamic simulations supported hypothesis that these hydrogen bond interactions are indeed sustained; and in this pose, UM_213 was very stable with an average RMSD of <1 Å (**Supplemental Figures 10-12**). Analogs were designed to assess the importance of the large hydrophobic cycloheptane group and to optimize hydrogen bonding interactions at the opposite end of the molecule, changing the indole group for an oxo-piperazine (**UM_259)** or an imidazolidine-2,4-dione (**UM_258**) **(Supplemental Figure 12)**. Docking of analogs supported the hypothesis that the chromenone-cycloheptanyl group fits in the binding pocket by the hinge and gatekeeper while increased hydrogen bonding interactions could be observed via Val93, Lys44 and Ser156 with additional bonds at Asp136 in the 4FG9 model for UM_259 (**Supplemental Figure 13, 14). UM_259** was experimentally determined to have a higher affinity for PNCK than the parent, UM_213.

**UM_195**, 5-[5-(naphthalen-1-yl)-1,3,4-oxadiazol-2-yl]sulfanylpyrazine-2-carbonitrile: While UM_195 was one of the most potent ligands as determined via the NanoBRET assay, modeling of UM_195 proved to be the most difficult. Kinase inhibitors classically bind with strong hydrogen bond interactions at the hinge. However, UM_195 docking poses in all 15 grids did not predict any such interactions. The naphthalene group was predicted to be deep in the binding pocket with hydrophobic interactions at the gatekeeper residue (**Figure 5**). Some hydrogen bonding to the oxadiazol, nitrile or pyrazine groups may occur with the catalytic lyisine. Molecular dynamics interactions shifted the compound into the binding pocket so that the nitrogen atoms on the oxadiazol interacted with Val93 of the hinge, leaving the nitrile group solvent exposed (**Supplemental Figures 15, 16)**. Analogs replacing the naphthalene group with heterocycles containing hydrogen bond donors and acceptors were virtually screened. Additionally, the nitrile group was replaced with various groups to improve hydrogen bonding interactions with Ser156, Lys44, Ser27 (**Supplemental Figure 17**). Docking of analogs, however, supported the initial binding hypothesis. Analogs tested (UM_242, UM_244, UM_245, UM_253, UM_254) represented these changes. **UM_254** proved to be the most potent analog (which substituted the terminal nitrile group with a methoxy-ester) with an IC50 of 2.68uM. However, molecular dynamics analysis of UM_254 in any of the three models did not predict stable binding in the active site.

**UM210**, 6-chloro-2-[(quinolin-2-ylsulfanyl)methyl]-3,4-dihydroquinazolin-4-one: Initial molecular modeling predicted that UM_210 binds to PNCK with hydrogen bonding interactions to the hinge residue, Val93, via the chlorine atom on the fused pyrimidine heterocycle (**Supplemental Figures 18, 19)**. It was considered more likely that the chlorine residue is solvent exposed and the heterocycle oxygen forming a hydrogen bond with either Lys44 or Val93. Indeed, molecular dynamics simulations of UM_210 in both the 4FG8 and 4FG9 models situated the compound in the active site with conserved hinge interactions at the pyrimidine scaffold, sustaining contacts with Val93 and Glu91. Analogs were selected for testing to assess the likely binding mode of the compound (**Supplemental Figure 20)**. For example, the quinoline ring was changed to an indazole or pyridine group. Additionally, the chlorine was removed to determine if this atom in this position contributed significantly to binding. Docking of the analogs, however, supported another binding hypothesis whereby the quinoline ring was situated deep in the binding pocket with the benzene ring extending into the gatekeeper region while the nitrogen on the quinoline participated in hydrogen bonding with hinge residues. Analogs tested, UM_278, UM_279 and UM_280 did not demonstrate significantly improved binding in NanoBRET assays.

**UM_037**,

1-(3-5H-[1,2,4]triazino[5,6-b]indol-3-ylsulfanylpropyl)-2,3-dihydro-1H-1,3-benzodiazol-2-one: UM_037 was the highest scoring compound in every docking model of PNCK. It was initially screened in ADP-KinaseGLO where it had minimal inhibitory activity. However, it showed significant binding in NanoBRET with an IC50 of 8.79 uM. UM_037 contains two fused heterocycle moieties, both of which are common structures among hinge-interacting kinase inhibitors. The benzimidazolinone is predicted to be situated towards the hinge with interactions between the nitrogen NH and carbonyl oxygen with the Val93 backbone. Molecular dynamics also suggested additional interactions of the benzimidazolinone with Ser156 while the triazino group had hydrogen bonds with Lys44 and Ser156 and the indole group interacted with Glu140 (**Supplemental Figures 21, 22**). Additionally, there is likely pi-stacking interactions with these groups with Phe158. One model did orient the molecule in the opposite direction, with the indole group making interactions at the hinge. To determine the proper pose, analogs were designed that altered the triazino/indol group, including adding or removing bulky groups or adding carbonyl groups to alter hydrogen bonding (**Supplemental Figure 23)**. Molecular modeling of analogs supported the initial binding hypothesis whereby substituting the triazino-indol group for more flexible groups which retained hydrogen bonding ability resulted in better docking results. Despite improvement in docking scores, the analogs were not significantly better inhibitors of PNCK *in vitro*.

## Discussion

Computational methods are increasingly being employed in the preclinical hit discovery and lead optimization process to assist in the identification, prioritization, and optimization of active kinase inhibitor chemotypes. With the wide-spread availability of high-resolution crystals structures, large-scale chemical databases, and a slew of virtual screening platforms, *in silico* methods have contributed to the discovery of many novel kinase inhibitors in recent years. For example, several studies have used virtual screening to discover inhibitors for EGFR[154], BCR-ABL[155], PI3K[156], AURKA[157] and PKC[158]. Even understudied kinases were able to be “ drugged” via the use of homology modeling and machine learning as in the case of VEGFR2 (for which no crystal structure exists)[159]. Thus, there is strong precedent for the successes of such computational pipelines to generate novel discoveries, which otherwise likely would have taken much longer and at significantly higher cost[1]. The goal of such computational screening is the identification of chemically tractable starting points to use for medicinal chemistry optimizations towards a potent molecular probe or preclinical drug candidate. Computer aided drug discovery can result in a 1000-fold enrichment of hit discovery compared to traditional screening methods. Whereby HTS is limited to hundreds of thousands to a million compounds at enormous cost, a virtual screening campaign can readily screen tens to hundreds of million and, more recently even billions of compounds to prioritize a small subset of compounds to test in biochemical screens, dramatically increasing the likelihood of obtaining starting points for lead optimization[157]. Typical hit rates from experimental HTS can range between 0.01% and 0.14%, while hit rates for prospective virtual screen can be as high as 50%[160]. Various rounds of enrichment, including applying certain structure filters (PAINS, lead-like, drug-like), machine-learning predictions, 3D shape overlay, molecular docking, and molecular dynamics, contribute to the improved success of computational drug discovery efforts. There are two major methodologies for computer aided drug discovery: structure-based methods and ligand-based methods[161]. Structure based methods are built upon the foundation of an experimentally derived, high resolution crystal structure of the target protein and active site. Ligand based methods, conversely, are built upon the foundation of a known ligand or ligands for a particular target. Due to the fact that this ambitious project sought to drug an understudied kinase, there exists no crystal structure and no known exogenous ligands to use a foundation. Thus, several methods from both structure-based and ligand-based approaches were combined to produce a pipeline capable of predicting first-in-class active PNCK inhibitors.

The process of this computational drug discovery project involves multiple essential components. First, a reliable and highly predictive homology model of the target protein, PNCK, needed to be built. As PNCK is an understudied kinase, there are no reported crystal structures of this protein. However, structures of homologous kinases were readily available to use as a structural template for building such a model. Multiple protein ensembles were evaluated to most accurately and broadly sample the kinase conformational and energetic space. Second, a library of ligands needed to be curated for use in a virtual screening campaign. The selection of compounds to use in a screen is one of the largest contributors to success in hit identification[160, 162]. Compound curation and selection determines the size of the screening library, the scaffold diversity, and quality of positive and negative data[162]. Various computational methods are employed to filter molecules for favorable factors that have been noted in various large-scale studies of “ drug-like” and “ lead-like” properties. Most notable, “ Rule of Five” [163] or “ Rule of Three” [164] filters select for compounds with desirable molecular weights, lipophilicity and hydrogen bond interactors. More sophisticated curation efforts involve the use of pharmacophore searching, similarity screening, and AI-assisted machine learning. While it is impossible to sample the entirety of the chemical space, the use of several diverse, commercially available databases is a reasonable starting point to identify active chemical structures as these can be purchased off-the-shelf and tested rapidly. Here we used two distinct machine learning algorithms to survey >7million compounds collectively in eMolecules, MolDB and Enamine compound libraries. Through initial rounds of filtering based on predicted probabilities of activities as well as physiochemical properties, the number of compounds chosen for use in molecular docking studies was greatly reduced from millions to thousands. Following molecular docking, compounds were scored, ranked and prioritized for further analysis. Compounds were clustered using topological features to identify chemotypes that would be enriched for potent binding and inhibition. Docking poses and binding hypotheses were evaluated manually for the top scoring compounds. Scaffolds were additionally manually curated based on medicinal chemistry considerations, evaluating for synthetic tractability and preliminary ADMET properties or liabilities. Extended molecular dynamic simulations were run for select protein-compound complexes to further refine predicted protein-ligand complexes and to study the energetics of binding. Top compounds from each cluster were purchased for testing in various binding assays or screens. Data from those screens informed the selection or design of analogs (virtually or synthetically) which allowed for further rounds of preliminary optimizations.

In conclusion, 1 top Scaffold was identified from Naïve-Bayesian classification followed by docking while the other 4 were identified through the nucleotide shape and color screen followed by docking. The docking scores could not be directly compared across the two methods (GLIDE and HYBRID) due to varying algorithms and there was no clear correlation between the scores, with each model ranking and prioritizing compounds differently. Few compounds scored highly in both models, making it difficult to draw generalized conclusions. As all virtual screening methods, docking scores should not be interpreted as relative binding free energies, but used as a tool for enrichment. UM_039 is the highest ranked kinase in campaign 1 and subsequently had the highest level of inhibitory activity in the ADP Glo assay. Conversely, UM_228 and UM_195, two of the most potent scaffolds in the NanoBRET assay, rank quite low when scored by GLIDE. For the HYBRID docking used in campaign 3, UM_195 is amongst the top scoring compounds and subsequently is a top inhibitory scaffold. The overall success of the entire campaign is therefore likely due to the combination of diverse methods and compound libraries, which can result in synergies.

PNCK is an understudied kinase for which there currently exists no published crystal structure and no known exogenous ligands. However, using a combination of homology modeling, machine learning and shape based virtual screening, we report the first-in-class series of hit compounds with inhibitory activity against PNCK. None of these compounds, although commercially available, have any target annotation data and thus have not been previously characterized as kinase inhibitors. The overall “ hit rate” of 27% is exceptionally high due to rounds of iterative enrichment using machine learning and shape-based scoring with molecular docking and molecular dynamics studies. Specifically, all the compounds discovered have IC50s in the low micromolar range and molecular weights under 400 Daltons, making them a good starting point for hit-to-lead optimization. Future goals are to evaluate the target specificity of these initial hit compounds in a kinome-wide screen, with particular focus on developing isoform specific inhibitors of PNCK (vs other CAMK proteins). The most selective scaffold(s) will then be chosen for lead optimization using medicinal chemistry. With preliminary SAR by catalog studies, we have improved the IC50s of UM_228, UM_213 and UM_195. UM_251 and UM_252 are both analogs of UM_228 with IC50s of 2.13 and 1.09 uM respectively-making them the most potent PNCK inhibitors in the series. Additional rounds of optimizations will be needed to optimize their activity while improving ADMET properties. In particular, we hope to replace all thioethers with oxygen isosteres and eliminate any furan rings, which are known to be metabolic liabilities. Once an isoform specific and potent chemical probe is generated, with an IC50 in the nanomolar range, we will move forward for testing in cells and in animal models. Previous work has been done to develop potent CAMK1D inhibitors. While the authors achieved specificity to CAMKs, they were unable to obtain isoform specificity.

## Acknowledgements

This work was supported by NIH grants U54HL127624 (BD2K LINCS Data Coordination and Integration Center, DCIC), U24TR002278 (Illuminating the Druggable Genome Resource Dissemination and Outreach Center, IDG-RDOC), U01LM012630 (BD2K, Enhancing the efficiency and effectiveness of digital curation for biomedical ‘big data’). The authors would like to acknowledge Ron Lampert and Sarah Simon for their editing of review of manuscript and creation of figures and tables.

## Supporting Information Captions

**Supplemental Figure 1: Molecular Dynamics (MD) Analysis of 3 PNCK Homology Models**. (**A-C**), Molecular Dynamic RMSD analysis of protein (C-alpha) and ligand fit on protein (4FG7, 4FG8, 4FG9). (**D-F**) Interaction Fraction of ATP to Residues in PNCK Active Site (4FG7, 4FG8, 4FG9).

**Supplemental Figure 2: SwissModel Ramachandran Plots for PNCK Homology Models** A) 4FG7 Homology Model, B) 4FG7 Original, C) 4FG8 Homology Model, D) 4FG8 Original, E) 4FG9 Homology Model, F) 4FG9 Original

**Supplemental Figure 3: ADP-Kinase GLO Results from Naïve Bayesian Classifier Campaign**.

**Supplemental Figure 4: UM_228 Binding Mode Hypothesis After MD, 4FG8 Model** A) 2D representation, B) 3D Representation C) 3D Surface Representation of Binding Pocket

**Supplemental Figure 5: Molecular Dynamics Analysis of UM_228**. A) Protein-Ligand RMSD plot of 500ns MD simulation as a function of time. B) Protein-Ligand Contact Interactions as Fraction of total MD simulation time. C) Protein-Ligand Contact plot as function of time. Darker lines indicate more contacts between the compound and residue.

**Supplemental Figure 6: Structure Activity Relationship (SAR) Table for Analogs of UM_228**.

**Supplemental Figure 7: 2D Binding Hypothesis of UM_228 Analogs**. A) UM_251, B) UM_252.

**Supplemental Figure 8: Molecular Dynamics Analysis of UM_251**. A) Protein-Ligand RMSD plot of 500ns MD simulation as a function of time. B) Protein-Ligand Contact Interactions as Fraction of total MD simulation time. C) Protein-Ligand Contact plot as function of time. Darker lines indicate more contacts between the compound and residue.

**Supplemental Figure 9: Molecular Dynamics Analysis of UM_252**. A) Protein-Ligand RMSD plot of 500ns MD simulation as a function of time. B) Protein-Ligand Contact Interactions as Fraction of total MD simulation time. C) Protein-Ligand Contact plot as function of time. Darker lines indicate more contacts between the compound and residue.

**Supplemental Figure 10: UM_213 Binding Mode Hypothesis After MD, 4FG9 Model** A) 2D representation, B) 3D Representation C) 3D Surface Representation of Binding Pocket

**Supplemental Figure 11: Molecular Dynamics Analysis of UM_213**. A) Protein-Ligand RMSD plot of 500ns MD simulation as a function of time. B) Protein-Ligand Contact Interactions as Fraction of total MD simulation time. C) Protein-Ligand Contact plot as function of time. Darker lines indicate more contacts between the compound and residue.

**Supplemental Figure 12: Structure Activity Relationship (SAR) Table for Analogs of UM_213**.

**Supplemental Figure 13: 2D Binding Hypothesis of UM_213 Analog, UM_259**

**Supplemental Figure 14: Molecular Dynamics Analysis of UM_259**. A) Protein-Ligand RMSD plot of 500ns MD simulation as a function of time. B) Protein-Ligand Contact Interactions as Fraction of total MD simulation time. C) Protein-Ligand Contact plot as function of time. Darker lines indicate more contacts between the compound and residue.

**Supplemental Figure 15: UM195 Binding Mode Hypothesis After MD, 4FG7 Model** A) 2D representation, B) 3D Representation C) 3D Surface Representation of Binding Pocket

**Supplemental Figure 16: Molecular Dynamics Analysis of UM_195**. A) Protein-Ligand RMSD plot of 500ns MD simulation as a function of time. B) Protein-Ligand Contact Interactions as Fraction of total MD simulation time. C) Protein-Ligand Contact plot as function of time. Darker lines indicate more contacts between the compound and residue.

**Supplemental Figure 17: Structure Activity Relationship (SAR) Table for Analogs of UM_195**.

**Supplemental Figure 18: UM210 Binding Mode Hypothesis After MD, 4FG9 Model** A) 2D representation, B) 3D Representation C) 3D Surface Representation of Binding Pocket

**Supplemental Figure 19: Molecular Dynamics Analysis of UM_210**. A) Protein-Ligand RMSD plot of 500ns MD simulation as a function of time. B) Protein-Ligand Contact Interactions as Fraction of total MD simulation time. C) Protein-Ligand Contact plot as function of time. Darker lines indicate more contacts between the compound and residue.

**Supplemental Figure 20: Structure Activity Relationship (SAR) Table for Analogs of UM_210**.

**Supplemental Figure 21: UM37 Binding Mode Hypothesis After MD, 4FG8 Model** A) 2D representation, B) 3D Representation C) 3D Surface Representation of Binding Pocket

**Supplemental Figure 22: Molecular Dynamics Analysis of UM_37**. A) Protein-Ligand RMSD plot of 500ns MD simulation as a function of time. B) Protein-Ligand Contact Interactions as Fraction of total MD simulation time. C) Protein-Ligand Contact plot as function of time. Darker lines indicate more contacts between the compound and residue.

**Supplemental Figure 23: Structure Activity Relationship (SAR) Table for Analogs of UM_37**.

